# Seasonal photoperiod cycling reduces inter-individual variability in μ-opioid receptor density in rats

**DOI:** 10.64898/2026.08.03.742441

**Authors:** Lihua Sun, Heidi Liljenbäck, Jenni Virta, Johan Rajander, Semi Helin, Emrah Yatkin, Jing Tang, Anne Roivanen

**Author notes:** Correspondence to: Lihua Sun, Turku PET Centre, University of Turku, Turku FI-20520, Finland.

## Abstract

**Rationale:** The μ-opioid receptor (MOR) is widely expressed across tissues and plays crucial roles in pain and stress responses, social behaviour, and immune regulation. Recent evidence indicates seasonal variation in *in vivo* MOR signalling; for example, short photoperiods are associated with reduced central MOR availability, increased MOR expression in brown adipose tissue (BAT), and strengthened brain–BAT interactions. However, despite this coupling with photoperiod, it remains unclear whether static daylength and dynamically changing photoperiods exert distinct effects, as adaptation to photoperiod transitions may itself induce stress-related modulation of the MOR system. Elucidating how seasonal, stress-related adaptations influence MOR signalling is essential for advancing our understanding of seasonal fluctuations in mood and stress regulation.

**Methods:** We compared rats housed under seasonal photoperiod cycling with those maintained under constant photoperiod conditions, using *ex vivo* radioligand binding to directly assess MOR density in central and peripheral tissues.

**Results:** Rats exposed to seasonal photoperiod cycling showed markedly reduced inter-individual variability in MOR density in both the brain (including the cerebellum and striatum) and peripheral tissues (adrenal glands), whereas no tissues exhibited substantially increased variability.

**Conclusions:** These findings demonstrate that seasonal photoperiod cycling stabilizes MOR dynamics at the population level, suggesting stress-related synchronization of MOR signalling. The findings deepen our understanding of seasonal effects on endogenous MOR signalling, and further underscore the role of seasonal light variation in modulating mood-related processes.

## Introduction

The μ-opioid receptor (MOR) is widely expressed across tissues, including the brain, brown adipose tissue (BAT), adrenal glands, and spleen (Sun et al., 2023; Wittert et al., 1996). Despite the well-established role of central MOR signalling in pain regulation, stress responses, and social behaviour (Becker et al., 2014; Lihua Sun et al., 2021), the functions of peripheral MOR signalling have largely been limited to local analgesic effects and immune regulation (Stein et al., 1993; Stein et al., 2001; Vadivelu, 2011). Recently, seasonal variation in photoperiod has been shown to modulate endogenous opioid signalling, as demonstrated in both clinical observations and preclinical simulation studies. In particular, *in vivo* MOR signalling, quantified using PET as receptor availability, exhibits a seasonal pattern in which both extremely long and short photoperiods are associated with reduced central MOR availability (L. Sun et al., 2021). In peripheral BAT, shorter photoperiods increase MOR availability (Sun et al., 2023) and enhance brain-BAT crosstalk (Sun et al., 2025). Together, these findings substantially advance our understanding of the neurochemical mechanisms underlying seasonal fluctuations in mood and physiology (Sun, 2025).

Despite the evident association between photoperiod and MOR signalling, it remains unclear whether static daylength and dynamically changing photoperiods exert differential effects. Adaptation to transitions in photoperiod may itself constitute a stressor, potentially leading to stress-related modulation of the MOR system. Clarifying how such seasonal adaptations influence MOR signalling is therefore critical for understanding seasonal variation in mood and stress regulation. To address this question, we compared rats exposed to seasonally cycling photoperiods with rats maintained under constant photoperiod conditions. MOR density was quantified using *ex vivo* radioligand binding. We further extended the analysis across multiple brain regions and peripheral tissues, including components of the adrenal–brain axis relevant to stress responses. We hypothesized that seasonal photoperiod cycling induces stress-related adaptations that modulate MOR signalling.

## Methods

### Animals

Sixteen rats were included in the *ex vivo* gamma counting analyses. Of these, seven were exposed to a seasonally varying photoperiod designed to mimic natural changes in daylength (Experimental group), whereas nine were maintained under a constant 12:12 h light–dark cycle (Control group). The Experimental group comprised five males and two females (age: 178.3 ± 8.0 days), whereas the Control group included five males and four females (age: 203.7 ± 9.9 days). Longitudinal PET imaging was conducted in a subset of animals, including two males from the Experimental group and two females from the Control group. All the procedures and protocols are in accordance with the EU Directive 2010/63/EU on the protection of animals used for scientific purposes, and are approved by the National Animal Experiment Board of Finland (license: 3116/04.10.07/2017 and 8648/2020).

### *Ex vivo* gamma counting

Twenty minutes after intravenous administration of [^11^C]carfentanil (40 MBq), rats were euthanized under isoflurane anaesthesia. Peripheral tissues and selected brain regions were rapidly excised, weighed, and measured for radioactivity using a gamma counter (Triathler 3 inch, Hidex). Radioactivity concentrations were expressed as percentage of injected radioactivity dose per gram of tissue (%ID/g).

### Statistical analysis

#### Analysis of variability

Differences in variability between the Control and Experimental groups were assessed for each region of interest (ROI) using the Brown–Forsythe test, a robust modification of Levene’s test based on deviations from the group median. This approach was selected to account for non-normal distributions and potential outliers in the data.

For each ROI, variability was quantified using standard deviation (SD) and variance. To facilitate interpretation of group differences, a variance ratio (Control/Experimental) was calculated, with values greater than 1 indicating higher variability in the Control group. To account for multiple comparisons across ROIs, p-values were adjusted using the Benjamini– Hochberg false discovery rate (FDR) procedure. All analyses were conducted in R (version 4.6.0) using the *car* package.

#### Pooled analysis of age, daylength, and sex effects

Separate generalized linear models (GLMs) were fitted for each ROI. In each model, %ID/g was modelled as a function of age, sex, and daylength. Regression coefficients, standard errors, p-values, and 95% confidence intervals (CI) were extracted for each predictor. False discovery rate (FDR) correction was applied using the Benjamini–Hochberg procedure, with correction performed separately for each predictor category (age, sex, and daylength).

## Results

### Between-group differences in variability

A significant difference in variability between groups was observed in the cerebellum, where the Control group exhibited substantially greater dispersion than the Experimental group (SD: 0.40 vs. 0.041; variance: 0.162 vs. 0.0017; F = 15.17, p = 0.0016, FDR-corrected p = 0.028). In the adrenal glands, variability was markedly higher in the Control group (SD: 15.3 vs. 0.64; variance: 233 vs. 0.41; F = 4.99, p = 0.042); however, this effect did not remain significant after FDR correction (FDR-corrected p = 0.36). Similarly, a trend toward increased variability in the Control group was observed in the striatum (SD: 1.60 vs. 0.37; variance: 2.55 vs. 0.13; F = 3.79, p = 0.072).

Across several additional regions, including BAT, muscle, neocortex, spleen, hippocampus, hypothalamus, and intestines, the Control group consistently showed higher variability than the Experimental group (variance ratios > 1); however, these differences were not statistically significant.

**Figure 1.**
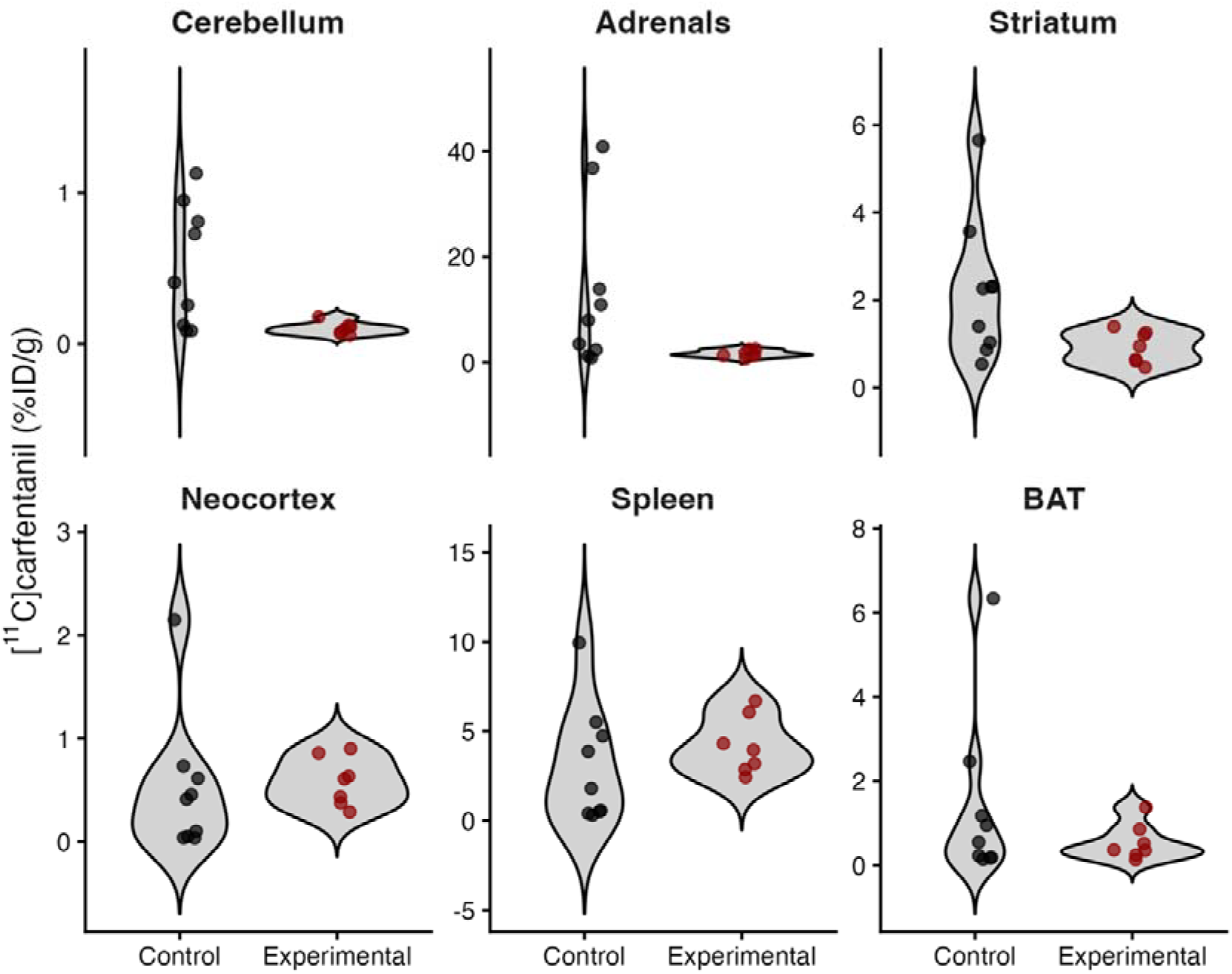
Seasonal photoperiod cycling synchronizes regional MOR density across animals. BAT = brown adipose tissue. Result for other regions are presented in Supplementary Figure S1.

In contrast, no evidence of greater variability in the Control group was observed in the liver or lungs, where variance was comparable or slightly higher in the Experimental group (variance ratios < 1). Other regions, including the kidney, pancreas, thalamus, white adipose tissue (WAT), and heart, showed only minor differences in dispersion between groups.

### Assessment of age, sex, and daylength effects

Animals in the Experimental group were younger and exposed to shorter daylengths at the time of sacrifice compared with those in the Control group (Wilcoxon rank-sum test: age, p = 5.25 × 10□□; daylength, p = 1.75 × 10□□).

After pooling the two groups, several regions showed nominal (uncorrected) associations with age, including the thalamus (β = 0.044, p = 0.011), neocortex (β = 0.028, p = 0.046), and WAT (β = −0.060, p = 0.029), however these associations did not remain significant after multiple-comparison correction (**Supplementary Table S1**). All other regions were non-significant (p > 0.05). For Daylength, nominal negative associations were observed in the thalamus (β = −0.261, p = 0.012) and neocortex (β = −0.171, p = 0.042), but none survived FDR correction.

In contrast, sex (male vs. female) showed robust effects. After FDR correction, males exhibited significantly lower values in the following regions: thalamus (β = −1.138, t = −4.52, p < 0.001, q = 0.012, 95% CI [−1.686, −0.590]); neocortex (β = −0.760, t = −3.55, p = 0.004, q = 0.026, 95% CI [−1.225, −0.294]); and spleen (β = −3.872, t = −3.49, p = 0.005, q = 0.026,5% CI [−6.293, −1.451]). No other sex differences survived correction, although nominal effects were observed in BAT, heart, and pancreas (p < 0.05, uncorrected). Full results are presented in **Supplementary Table S1**.

## Discussion

Our data show that the cycling photoperiod stabilizes MOR density in the cerebellum, adrenal glands, and striatum across animals. This may reflect stress-related synchronization of MOR signalling across individuals. These findings advance our understanding of seasonal variation in MOR signalling dynamics by introdeucing a novel dimension of seasonal effects, extending the regional scope to both central and peripheral tissues, and providing more direct evidence of receptor density changes. Overall, the findings indicate that seasonal photoperiod changes exert complex effects on the MOR system, further highlighting the role of MOR signalling in seasonal modulation of mood and stress responses.

MOR signalling plays an important role in regulating the hypothalamic–pituitary–adrenal (HPA) axis, which is central to the physiological response to stress. MORs are expressed in the adrenal cortex, and their activation can inhibit the secretion of adrenocorticotropic hormone and cortisol (Ducat et al., 2013; Krazinski et al., 2011). In our previous work, cycling photoperiod acted as an environmental stressor and was associated with increased serum corticosterone levels (L. Sun et al., 2021). In the present study, synchronization of adrenal MOR density across animals may therefore reflect a shared state of stress adaptation induced by photoperiod cycling.

MORs are expressed in cerebellar Purkinje cell in rats, where they may modulate excitatory input critical for cerebellar output and motor coordination (Mrkusich et al., 2004). In previous PET studies, the cerebellum was often been used as a reference region due to its relatively low MOR expression, a finding further confirmed in the present study using *ex vivo* radioligand binding. Nevertheless, we observed pronounced synchronization of MOR expression patterns in the cerebellum following seasonal photoperiod cycling. These findings suggest that chronic photoperiodic manipulation may induce coordinated neuroadaptive changes in cerebellar opioid signaling. Such alterations could influence motor-related processing and behavioral responsiveness, supporting the emerging view that the cerebellum may play a more active role in stress adaptation and affective regulation than previously appreciated (Gheorghe et al., 2018).

In the pooled analysis, Age and Daylength exhibited opposite effects, largely reflecting the study design whereby rats in the Experimental group were younger and temporarily exposed to shorter photoperiods. Importantly, these associations were not observed in the same regions where Group significantly affected the variability of the measures, suggesting that they are unlikely to account for the primary group effects. In addition, male rats exhibited lower MOR density across multiple brain and peripheral regions. These findings are consistent with previous reports indicating greater sensitivity to opioid analgesia in females (Craft, 2003).

In the present study, *ex vivo* gamma counting following intravenous administration of [^11^C]carfentanil provided a direct measure of tissue radioligand uptake (%ID/g). Compared with *in vivo* PET imaging, this approach reduces sources of measurement variability, including motion artefacts, image reconstruction noise, and partial volume effects. Although this method does not quantify absolute receptor density in a strict pharmacological sense, radioligand uptake likely to predominantly reflect MOR density because of the high selectivity and affinity of carfentanil for MORs.

### Limitations

Despite the relatively small sample size, the observed effects were robust and statistically significant. Nevertheless, several limitations should be acknowledged. In particular, the between-group imbalance may complicate interpretation of the findings, as animals in the Experimental group were younger and exposed to shorter photoperiod conditions at the time of sacrifice. However, effects of Age and Daylength were localized to brain regions distinct from those exhibiting the principal group differences in MOR density variability. These observations support the interpretation that the reported findings are primarily attributable to chronic photoperiod cycling, likely reflecting stress-related neurobiological adaptations induced by repeated photoperiodic transitions.

### Conclusions

These findings demonstrate that seasonal photoperiod cycling stabilizes MOR dynamics at the population level, indicating stress-related synchronization of MOR signaling across tissues implicated in affective and behavioral regulation. Given the established role of the MOR system in stress responsivity and social behavior, the present results further support the notion that seasonal light variation exerts broad modulatory effects on this neurochemical pathway. The observed synchronization of MOR density may reflect an adaptive neurobiological response to repeated environmental stressor. Collectively, these findings extend current understanding of seasonal influences on the endogenous opioid system and further highlight photoperiodic variation as a potential regulator of mood-related processes and stress adaptation.

## Author Contributions

Contributions of the authors include conception and design (LS, AR); data collection (HL, JV, JR, SH), analysis and interpretation of data (LS); drafting of the manuscript (LS); revising and comment (JR, SH, EY, JT, AR); and all authors approved the final manuscript for submission.

## Acknowledgement

The study is supported by Turku Collegium for Science, Medicine and Technology, University of Turku (LS) & Fudan University affiliated Huashan Hospital Starting Fund (#30302171001; LS).

