## Supplementary for "Seasonal photoperiod cycling reduces inter-individual variability in μ-opioid receptor density in rats"

Lihua Sun et al.

**Supplementary Results**

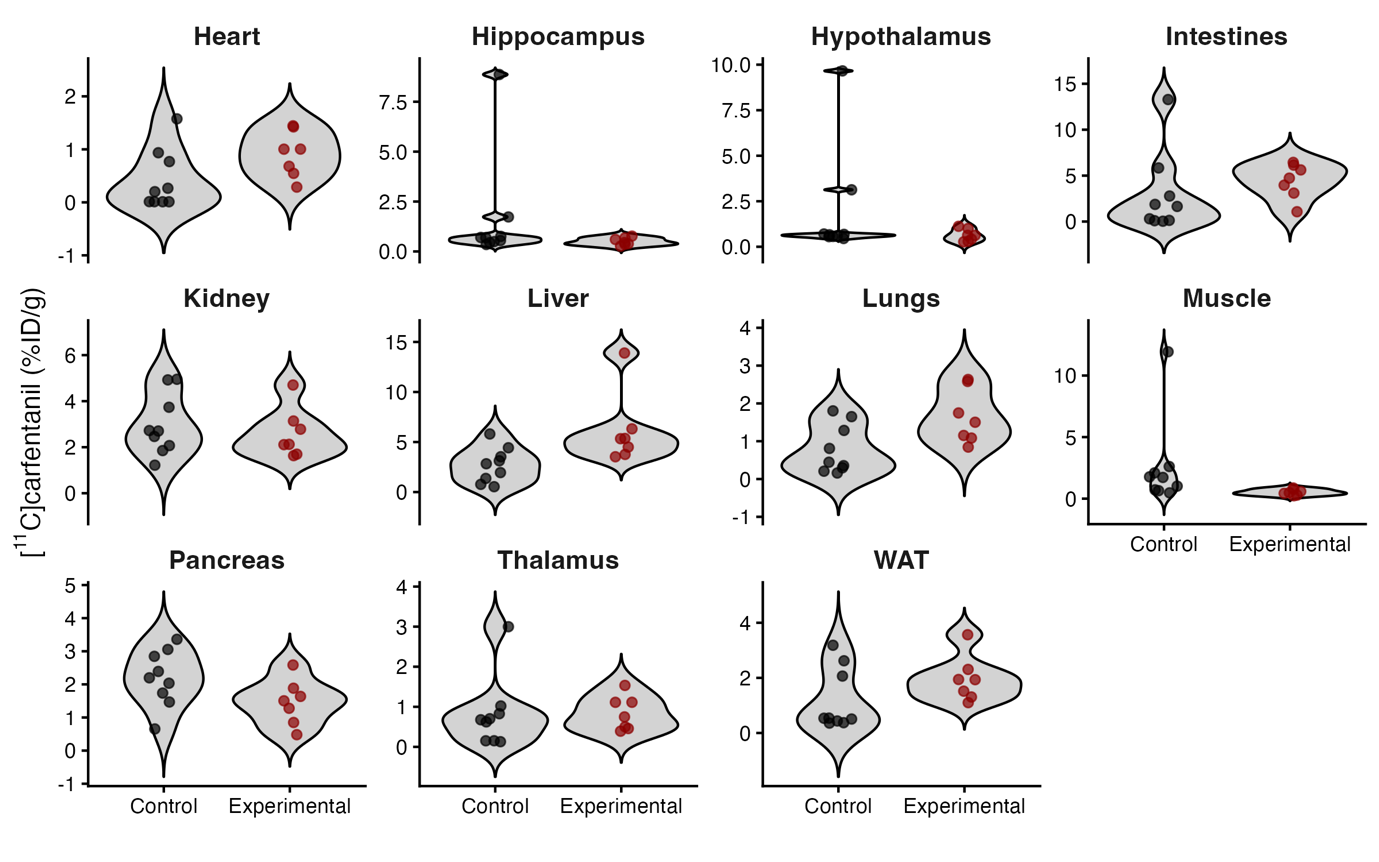

***Supplementary Figure S1.*** *Seasonal photoperiod cycling stabilizes MOR density across animals in other regions*

| ***Supplementary Table S1.*** *Effect of age, sex (male vs. female), and daylength on MOR density.*  **=p<0.05; **=p<0.01; ***=p<0.001.* | | | | | | | | | |
| --- | --- | --- | --- | --- | --- | --- | --- | --- | --- |
| **Predictor** | **ROI** | **Beta** | **SE** | **t value** | **p** | **Adj. p** | **95% CI**  **lower** | **95% CI**  **upper** | **FDR** |
| Age | Adrenals | 0.113 | 0.41 | 0.276 | 0.787 | 0.892 | -0.78 | 1.007 | No |
| Age | BAT | 0.014 | 0.045 | 0.31 | 0.762 | 0.892 | -0.085 | 0.113 | No |
| Age | Cerebellum | 0.02 | 0.009 | 2.082 | 0.059 | 0.253 | -0.001 | 0.04 | No |
| Age | Heart | 0.005 | 0.014 | 0.353 | 0.73 | 0.892 | -0.025 | 0.035 | No |
| Age | Hippocampus | 0.012 | 0.066 | 0.181 | 0.859 | 0.913 | -0.132 | 0.156 | No |
| Age | Hypothalamus | 0.035 | 0.07 | 0.499 | 0.627 | 0.888 | -0.118 | 0.188 | No |
| Age | Intestines | -0.121 | 0.101 | -1.193 | 0.256 | 0.597 | -0.341 | 0.1 | No |
| Age | Kidney | 0.041 | 0.038 | 1.069 | 0.306 | 0.597 | -0.042 | 0.124 | No |
| Age | Liver | -0.096 | 0.092 | -1.046 | 0.316 | 0.597 | -0.296 | 0.104 | No |
| Age | Lungs | -0.01 | 0.018 | -0.567 | 0.581 | 0.888 | -0.049 | 0.029 | No |
| Age | Muscle | 0.006 | 0.092 | 0.065 | 0.95 | 0.95 | -0.195 | 0.207 | No |
| Age | Neocortex | 0.028 | 0.012 | 2.22 | 0.046* | 0.253 | 0.001 | 0.055 | No |
| Age | Pancreas | 0.034 | 0.024 | 1.426 | 0.179 | 0.508 | -0.018 | 0.086 | No |
| Age | Spleen | 0.057 | 0.065 | 0.879 | 0.397 | 0.674 | -0.084 | 0.198 | No |
| Age | Striatum | 0.061 | 0.041 | 1.492 | 0.161 | 0.508 | -0.028 | 0.151 | No |
| Age | Thalamus | 0.044 | 0.015 | 3.02 | 0.011* | 0.181 | 0.012 | 0.076 | No |
| Age | WAT | -0.06 | 0.024 | -2.49 | 0.029* | 0.242 | -0.113 | -0.008 | No |
| Daylength | Adrenals | 1.49 | 2.49 | 0.599 | 0.561 | 0.881 | -3.934 | 6.914 | No |
| Daylength | BAT | 0.061 | 0.275 | 0.221 | 0.829 | 0.881 | -0.539 | 0.661 | No |
| Daylength | Cerebellum | -0.029 | 0.057 | -0.512 | 0.618 | 0.881 | -0.155 | 0.096 | No |
| Daylength | Heart | -0.131 | 0.084 | -1.566 | 0.143 | 0.665 | -0.313 | 0.051 | No |
| Daylength | Hippocampus | 0.137 | 0.402 | 0.34 | 0.74 | 0.881 | -0.739 | 1.012 | No |
| Daylength | Hypothalamus | 0.031 | 0.426 | 0.072 | 0.944 | 0.944 | -0.897 | 0.959 | No |
| Daylength | Intestines | 0.34 | 0.614 | 0.554 | 0.59 | 0.881 | -0.997 | 1.678 | No |
| Daylength | Kidney | -0.169 | 0.232 | -0.731 | 0.479 | 0.881 | -0.675 | 0.336 | No |
| Daylength | Liver | -0.139 | 0.558 | -0.249 | 0.807 | 0.881 | -1.354 | 1.076 | No |
| Daylength | Lungs | -0.149 | 0.108 | -1.371 | 0.196 | 0.665 | -0.385 | 0.088 | No |
| Daylength | Muscle | 0.34 | 0.561 | 0.606 | 0.556 | 0.881 | -0.881 | 1.561 | No |
| Daylength | Neocortex | -0.171 | 0.075 | -2.271 | 0.042* | 0.36 | -0.336 | -0.007 | No |
| Daylength | Pancreas | -0.055 | 0.144 | -0.381 | 0.71 | 0.881 | -0.369 | 0.259 | No |
| Daylength | Spleen | -0.582 | 0.392 | -1.484 | 0.164 | 0.665 | -1.437 | 0.273 | No |
| Daylength | Striatum | -0.106 | 0.249 | -0.425 | 0.679 | 0.881 | -0.649 | 0.437 | No |
| Daylength | Thalamus | -0.261 | 0.089 | -2.937 | 0.012* | 0.211 | -0.454 | -0.067 | No |
| Daylength | WAT | 0.142 | 0.147 | 0.968 | 0.352 | 0.881 | -0.178 | 0.463 | No |
| Sex | Adrenals | -1.693 | 7.051 | -0.24 | 0.814 | 0.814 | -17.06 | 13.669 | No |
| Sex | BAT | -1.745 | 0.78 | -2.237 | 0.045 | 0.153 | -3.444 | -0.045 | No |
| Sex | Cerebellum | -0.04 | 0.163 | -0.248 | 0.808 | 0.814 | -0.395 | 0.314 | No |
| Sex | Heart | -0.595 | 0.237 | -2.513 | 0.0273 | 0.116 | -1.112 | -0.079 | No |
| Sex | Hippocampus | -1.767 | 1.138 | -1.552 | 0.147 | 0.249 | -4.247 | 0.713 | No |
| Sex | Hypothalamus | -2.402 | 1.206 | -1.991 | 0.0697 | 0.169 | -5.031 | 0.227 | No |
| Sex | Intestines | -2.515 | 1.739 | -1.447 | 0.174 | 0.268 | -6.303 | 1.273 | No |
| Sex | Kidney | -1.076 | 0.657 | -1.638 | 0.127 | 0.24 | -2.507 | 0.355 | No |
| Sex | Liver | 0.465 | 1.579 | 0.294 | 0.774 | 0.814 | -2.976 | 3.906 | No |
| Sex | Lungs | -0.581 | 0.307 | -1.892 | 0.0829 | 0.176 | -1.25 | 0.088 | No |
| Sex | Muscle | -1.472 | 1.587 | -0.928 | 0.372 | 0.486 | -4.931 | 1.986 | No |
| Sex | Neocortex | -0.76 | 0.214 | -3.554 | 0.004** | 0.0255 | -1.225 | -0.294 | **Yes** |
| Sex | Pancreas | -0.866 | 0.409 | -2.118 | 0.056 | 0.158 | -1.756 | 0.025 | No |
| Sex | Spleen | -3.872 | 1.111 | -3.485 | 0.0045** | 0.0255 | -6.293 | -1.451 | **Yes** |
| Sex | Striatum | -0.574 | 0.706 | -0.813 | 0.432 | 0.525 | -2.113 | 0.965 | No |
| Sex | Thalamus | -1.138 | 0.252 | -4.523 | 0.0007*** | 0.0119 | -1.686 | -0.59 | **Yes** |
| Sex | WAT | -0.557 | 0.417 | -1.337 | 0.206 | 0.292 | -1.465 | 0.351 | No |
